# Intensity warping for multisite MRI harmonization

**DOI:** 10.1101/679357

**Authors:** J Wrobel, ML Martin, R Bakshi, PA Calabresi, M Elliot, D Raolf, RC Gur, RE Gur, RG Henry, G Nair, J Oh, N Papinutto, D Pelletier, DS Reich, W Rooney, TD Satterthwaite, W Stern, K Prabhakaran, N Sicotte, RT Shinohara, J Goldsmith, on behalf of the NAIMS Cooperative

## Abstract

In multisite neuroimaging studies there is often unwanted technical variation across scanners and sites. These “scanner effects” can hinder detection of biological features of interest, produce inconsistent results, and lead to spurious associations. We assess scanner effects in two brain magnetic resonance imaging (MRI) studies where subjects were measured on multiple scanners within a short time frame, so that one could assume any differences between images were due to technical rather than biological effects. We propose *mica* (**m**ultisite **i**mage harmonization by **C**DF **a**lignment), a tool to harmonize images taken on different scanners by identifying and removing within-subject scanner effects. Our goals in the present study were to (1) establish a method that removes scanner effects by leveraging multiple scans collected on the same subject, and, building on this, (2) develop a technique to quantify scanner effects in large multisite trials so these can be reduced as a preprocessing step. We found that unharmonized images were highly variable across site and scanner type, and our method effectively removed this variability by warping intensity distributions. We further studied the ability to predict intensity harmonization results for a scan taken on an existing subject at a new site using cross-validation.

## 1 Introduction

Medical imaging has become an established practice in clinical studies and medical research, leading to situations where images must be compared across site locations, scanners, or scanner types. Upgrades in scanner technology within a site may render old data not comparable to data collected on a newer machine, and this presents challenges in studies where acquisition techniques change over time. Multisite studies have become common as well; examples include large neuroimaging studies such as the Alzheimer’s Disease Neuroimaging Initiative (Mueller et al., 2005) and the Human Connectome Project (Van Essen et al., 2013), as well as targeted clinical trials studying multiple sclerosis (MS) interventions such as Kappos et al. (2006) and Hauser et al. (2017).

Measurement across multiple sites and scanners introduces unwanted technical variability in the images (Schnack et al., 2004). Going forward we will refer to technical artifacts introduced across either sites or scanners as “scanner effects.” Scanner effects in imaging studies can reduce power to detect true differences across images and distort downstream measurements of regional volumes, brain lesions, and other biological features of interest (Schnack et al., 2010; Jovicich et al., 2013; Cannon et al., 2014; Keshavan et al., 2016; Schwartz et al., 2019). In structural magnetic resonance imaging (MRI) studies, detection of scanner effects is particularly challenging because images are collected in arbitrary units of voxel intensity; as a result, raw MRI intensities are often not comparable across study visits even within the same subject and scanner. We refer to unwanted technical variability within the same scanner and subject that are due to arbitrary unit intensity values as “intensity unit effects.” Though often conflated, intensity unit effects and scanner effects are distinct sources of unwanted technical variation and should be treated separately. We refer to methods intended to address intensity unit effects as “normalization” methods to distinguish them from methods intended to reduce scanner effects, which we term “harmonization” methods. Both scanner effects and unit effects are present in multisite MRI studies, and in practice they can be challenging to separate.

Scanner effects can be due to differences in scanner hardware, scanner software, scan acquisition protocol, or other unknown sources. When present, images collected at different sites may have systematically different distributions of intensity values. For example, Shinohara et al. (2017) showed substantial differences in volumetrics across sites and scanner types even for a single, biologically stable subject measured under standardized protocols at the same field strength on platforms produced by the same vendor. The left panel of Figure 1 shows histograms of intensity values for this single subject, who was scanned twice at each of seven sites across the U.S. Large scanner effects are evident; smaller but visible differences within site show that intensity unit effects are present as well. In subsequent analyses, scanner effects produced inconsistent measurements of MS lesion volume both when lesions were segmented manually or by a variety of automated software pipelines (Shinohara et al., 2017).

**Figure 1:**
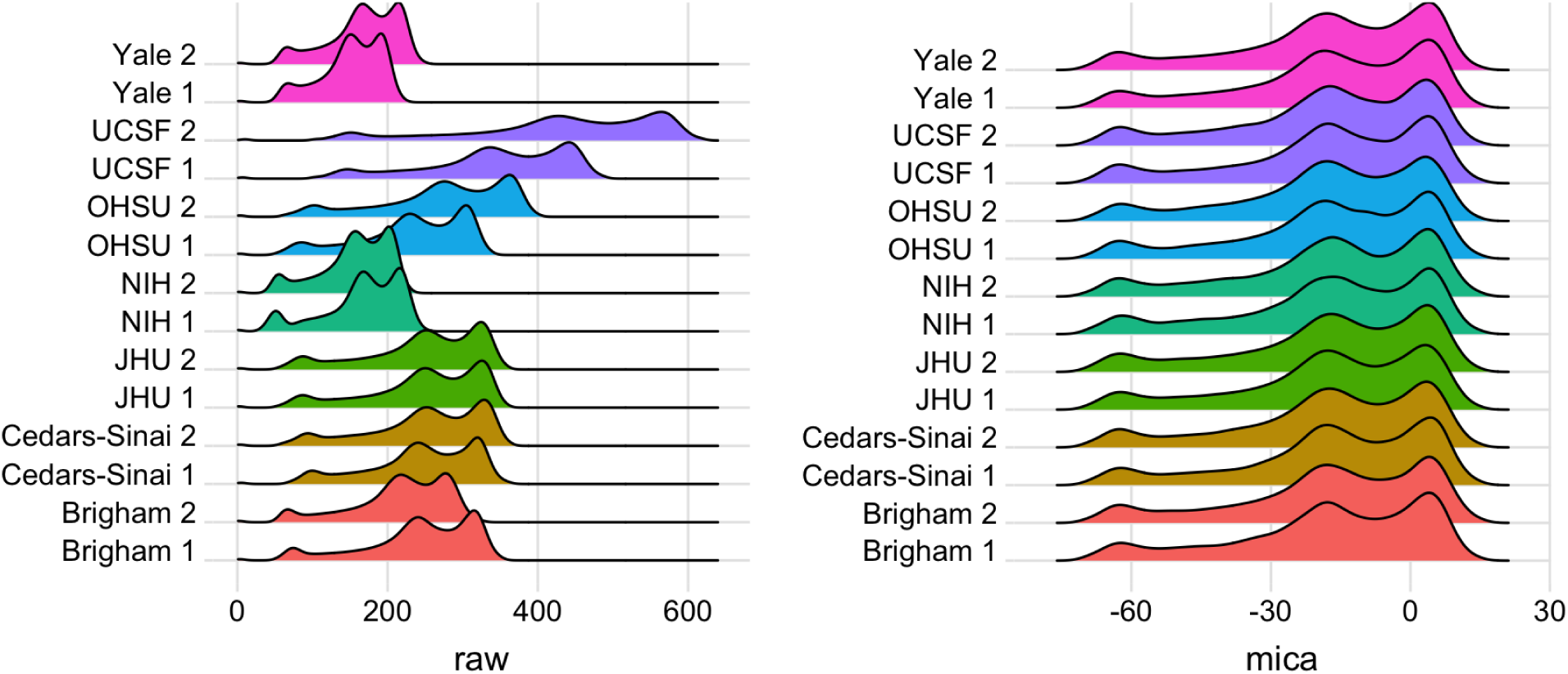
Histograms of voxel intensities for scan-rescan data across seven sites in the NAIMS pilot study: Brigham and Women’s Hospital (Brigham), Cedars-Sinai, Johns Hopkins University (JHU), National Institutes of Health (NIH), Oregon Health & Sciences University (OHSU), University of California San Francisco (UCSF), and Yale University (Yale). Left panel shows raw voxel intensities; right panel shows densities after *mica* harmonization and White Stripe normalization. At each site two scans were collected; a 1 or 2 after site name indicates the first or second scan, respectively.

The issue of arbitrary units has long been recognized and is the subject of a large literature on intensity normalization (Nyúl et al., 1999; Shinohara et al., 2011, 2014; Ghassemi et al., 2015). Intensity normalization methods facilitate comparability across subjects measured on the same scanner and standardize voxel intensity values; for a review of several methods see Shah et al. (2011). Histogram matching is an early approach that aligns densities of voxel intensities to quantiles of an image template constructed from several control subjects. Though popular, histogram matching often fails to preserve biological characteristics of individual scans and removes useful information regarding variation among subjects. Shinohara et al. (2014) formalized the principles of image normalization and introduced the White Stripe method. White Stripe normalizes images using patches of normal appearing white matter (NAWM), so that rescaled intensity values are biologically interpretable as units of NAWM. White Stripe can effectively normalize white matter across subjects and is a useful preprocessing step for automated lesion segmentation in MS (Sweeney et al., 2013a,b; Valcarcel et al., 2018), but technical variability can remain in the gray matter.

Unlike intensity normalization methods, which target intensity unit effects, harmonization methods aim to reduce scanner effects so that downstream analyses are more comparable across sites and scanners (Fortin et al., 2017; Yu et al., 2018). Fortin et al. (2018) described a voxel-wise regression method, based on tools from genomics, that harmonizes cortical thickness measurements from MRI scans. This method succeeds in removing scanner effects for measurements extracted from each image; in contrast, our goal in the present study was to develop an effective harmonization method that can be applied to the entire brain. Similar tools from genomics are used to correct for scanner effects in multisite diffusion tensor imaging data (Fortin et al., 2017) and multisite functional MRI data (Yu et al., 2018). However, these harmonization methods require spatial registration to a population template, which can lower image resolution and make it challenging to detect important disease features such as MS lesions. Ideally, an all-purpose harmonization method would remove scanner effects from the whole brain without requiring that all subjects be spatially registered to the same template image.

In the past, “normalization” has been used to simultaneously address the problems we characterize as unit and scanner effects, although these are more correctly viewed as distinct problems. As a result, intensity normalization techniques such as histogram matching and White Stripe are often used to address harmonization issues (Schnack et al., 2004; Shinohara et al., 2014; Fortin et al., 2016). Unlike harmonization techniques mentioned previously, these normalization techniques can be applied to the whole brain, do not require spatial registration, and reduce intensity unit effects. When scanner effects are due to the same voxel intensity transformations used to reduce unit effects, the normalization techniques will reduce scanner effects as well. However, often they fail to reduce much of the variability across sites, especially when large nonlinear scanner effects are present. Additionally, histogram matching normalizes voxel intensities across images at the cost of removing biological variability across subjects which can distort structures and mask inter-subject differences of interest.

Here, we introduce a new image intensity harmonization framework for multisite studies. We use data in which a subject was scanned on multiple scanners closely enough in time that any image differences can be attributed to differences across acquisition platforms (scanner effects) rather than biological effects. Our objectives in this study were to (1) establish a method that removes scanner effects by leveraging multiple scans collected on the same subject, and, building on this, (2) develop a technique to estimate scanner effects in large multisite trials so these can be reduced with preprocessing steps. The first objective establishes a framework for understanding harmonization, and the second relates to the practical use of this framework in multisite studies. We propose **m**ultisite **i**mage harmonization by **C**DF **a**lignment (*mica*), which harmonizes images by aligning cumulative distribution functions (CDFs) of voxel intensities. Our approach estimates nonlinear, monotonically increasing transformations of the voxel intensity values in one scan such that the resulting intensity CDF perfectly matches the intensity CDF from a second (“target”) scan. CDFs can be perfectly aligned using standard approaches to curve registration in the functional data analysis literature (Srivastava et al., 2011; Tucker et al., 2013; Wrobel et al., 2018). Although these intensity transformations, called warping functions, are defined using CDFs, they can be applied to voxel-level intensity values to produce a harmonized image. For a subject measured on different scanners in close succession, this allows us to identify and remove scanner effects; mappings established in this way can be used to reduce the impact of scanner effects in multisite studies.

We outline our harmonization approach using two data sets with distinct but related problems. The North American Imaging in Multiple Sclerosis (NAIMS) pilot study (Shinohara et al., 2017; Dworkin et al., 2018; Oh et al., 2018; Papinutto et al., 2018; Schwartz et al., 2019) found large scanner effects in a single subject with biologically stable MS, and we use these data to show that *mica* can reduce technical variability across sites while preserving the ability to detect MS lesions. A second study, which we refer to as the trio2prisma study, scanned ten healthy subjects on two different machines and found systematic nonlinear differences between the scanners. We used *mica* to harmonize images from the first scanner so that they are comparable to images collected on the second scanner; this demonstrates how our method can be used to create a mapping between scanners, and that scanner effects can be removed when data are available from both scanners for all subjects. Since scan-rescan data are often only available for a subset of study subjects, we also employed a leave-one-scan-out cross-validation approach to assess the utility of our harmonization method in this common setting. For both studies, we used *mica* to understand and, to the extent possible, remove scanner variability. We paired our method with White Stripe to remove intensity unit effects as well as scanner effects, though other intensity normalization methods could be used instead.

In the next section, we describe our data and the *mica* methodology. We then present the results of our technique in different settings, followed by a discussion.

### 2 Materials and Methods

### 2.1 Data and processing

#### 2.1.1 NAIMS dataset

The NAIMS steering committee developed a brain MRI protocol relevant to MS lesion quantification (Shinohara et al., 2017). Using this protocol, two scans were collected at each of seven sites across the United States on a 45-year-old man with clinically stable relapsing-remitting MS. All scans were performed on 3T Siemens scanners (four Skyra, two TimTrio, and one Verio). At each site, scan-rescan imaging was performed on the same day, with the subject exiting the machine between scans. The participant was also assessed at the beginning and end of the study on the same scanner to confirm disease stability by clinical and MRI measures.

Each image was bias-corrected using the N4 inhomogeneity correction algorithm (Tustison et al., 2010), then brain extraction was performed using the FSL BET skull-stripping algorithm (Smith, 2002). After performing *mica* harmonization as described in Section (2.2), T1-weighted (T1-w) and fluid attenuated inversion recovery (FLAIR) images were White Stripe normalized (Shinohara, R T and Muschelli, J, 2018) to remove intensity unit effects and enable automated MS lesion detection using the MIMoSA (Valcarcel et al., 2018) software pipeline.

#### 2.1.2 trio2prisma dataset

The trio2prisma data were collected from ten healthy subjects ages 19 to 29 at the University of Pennsylvania. For each subject, brain MRI scans were obtained on both a Siemens Trio machine and a Prisma scanner. Scans were performed between 2 and 11 days apart for each subject (mean 4.2 days), a time window in which we expect no significant structural changes in the brain. We focused on T1-w images for the trio2prisma data, though our method can be applied to other modalities as well. Images were bias-corrected, skull-stripped, and White Stripe normalized using the same algorithms described for the NAIMS data. Because normalization methods have often been used for harmonization in the past, we compared *mica* to White Stripe and histogram matching normalization. To assess method performance on this data, we compared white and gray matter segmentations for *mica*-harmonized images to White Stripe and histogram matching normalized images. All white and gray matter segmentations were obtained using multi-atlas Joint Label Fusion (Wang et al., 2013).

### 2.2 Methodology

Our framework for image harmonization uses non-linear transformations of image intensity values to remove scanner effects. The transformations were calculated by aligning distribution functions of intensity values. For a particular imaging modality (for example, T1-w), *Y*_*ijk*_(*v*) represents the intensity at a given voxel *v* for scan *j* of subject *i* measured at site *k*. Then *f*_*ijk*_(*x*) and *F*_*ijk*_(*x*) represent the probability density function (PDF) and CDF, respectively, for the voxel intensities of image *Y*_*ijk*_ measured over intensities *x*. Within each subject we assumed variability in voxel intensities across visits *j* and sites *k* is due to scanner and intensity unit effects rather than biological change, and that non-biological differences could be removed by aligning all CDFs for the *i*^*th*^ subject to a subject-specific “template CDF,” *F*_*it*_(*x*), for template *t*; template choices for our motivating studies are described below.

For image *Y*_*ijk*_ we estimate the nonlinear monotonic transformation of the intensity values, or *warping function*, 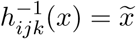, which aligns the CDF *F*_*ijk*_(*x*) to its template via

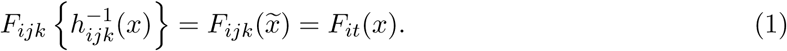

After alignment, the CDF of the original images becomes identical to the CDF of the template. For this reason, we use the notation *F*_*it*_(*x*) to represent the *mica*-harmonized CDF as well as the template for alignment. We further denote 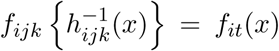 and *Y*_*it*_(*v*) to be the *mica*-harmonized PDFs and images, respectively. The aligned PDFs, *f*_*it*_(*x*), can be recovered from CDFs by differentiation. The warping functions 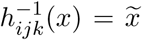 define a new intensity value, 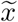, for each original intensity value in *x*. Since each *Y*_*ijk*_(*υ*) is a voxel intensity in *x*, harmonized images *Y*_*it*_ take values in 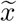 and are obtained by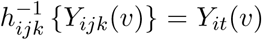. Figure (2) shows a schematic of this process: images were bias corrected and skull-stripped, voxel-intensities were converted to CDFs, CDFs were aligned, and warping functions from CDF alignment were used to generate harmonized images.

**Figure 2:**
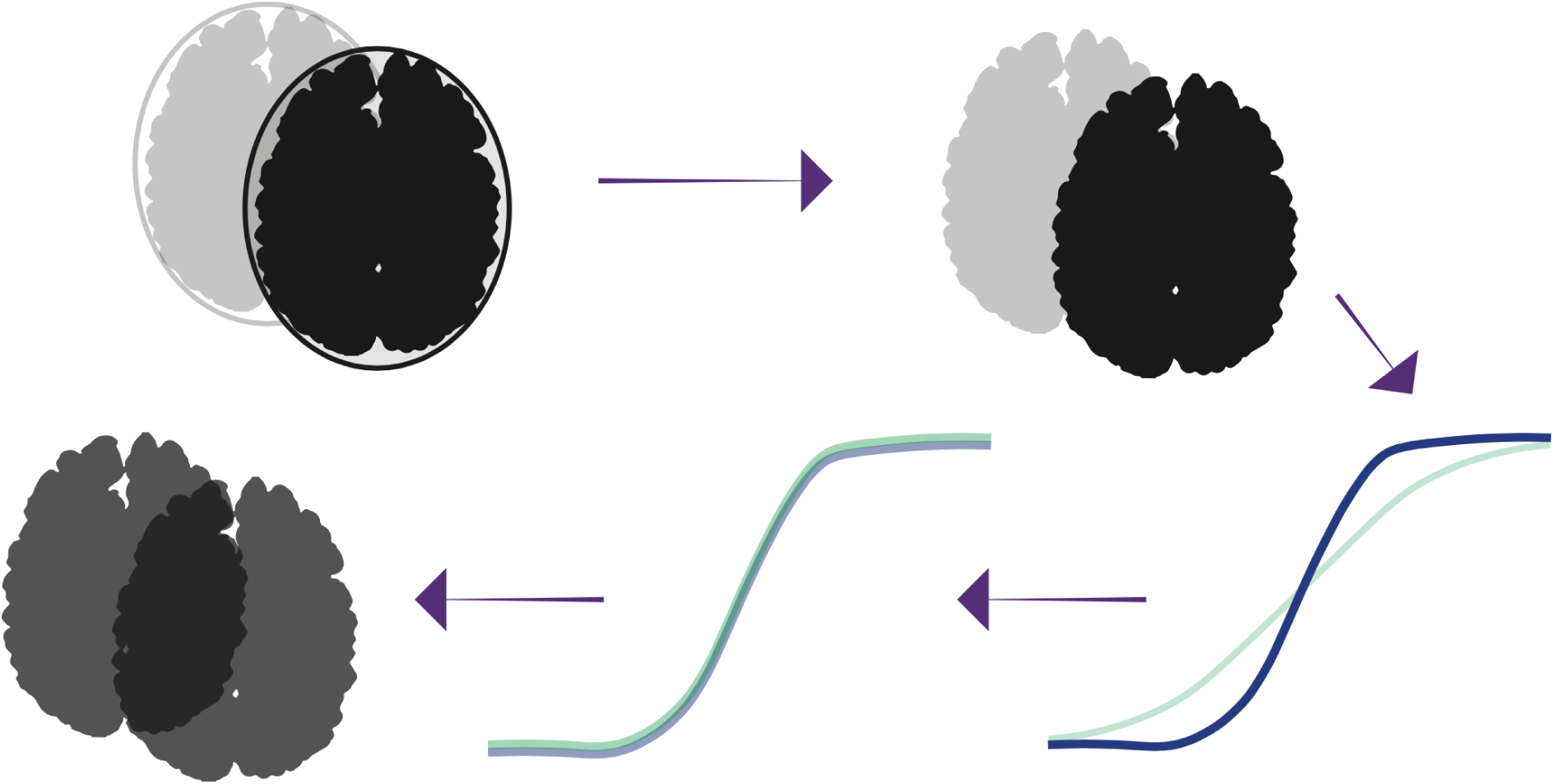
Harmonization pipeline. Raw images are N4 bias-corrected, skull-stripped, voxel intensities are converted to CDFs, CDFs are aligned by warping intensity values. The transformation of intensity values that produces this alignment is called a warping function, and the nonlinear transformation is applied to the raw images to produce harmonized images.

Given this framework for quantifying scanner effects, we now address objectives (1) and (2) stated in Section (1). Our first objective, to establish a method that removes scanner effects, is illustrated using both the NAIMS and the trio2prisma data. For NAIMS data, we obtained empirical CDFs of T1-w and FLAIR images from the NAIMS dataset. Within an imaging modality, each CDF is given by *F*_*ijk*_(*x*), *i* = 1, *j* ∈ {1, 2}, *k* ∈ {1, …, 7}. We used the Karcher mean as the common template *F*_*it*_(*x*) to which all CDFs within a modality are aligned, though in principle other templates could be used. For the trio2prisma data, we obtained empirical CDFs of T1-w images. Each CDF is given by *F*_*ijk*_(*x*), *i* ∈ {1, …, 10}, *j* = 1, *k* ∈ {Trio, Prisma}. For each subject, we used the CDF from the Prisma image, *F*_*i*Prisma_(*x*), as the template to which we align the CDF from the Trio image, *F*_*i*Trio_(*x*). Functions from the `fdasrvf` R package (Tucker, 2017) were used to perform alignment.

Our second objective was to develop a technique to estimate scanner effects in large multisite trials; to illustrate this, we used warping functions from the trio2prisma data. In such studies, most subjects are only measured on a single scanner. At best, only a subset of subjects will have scans collected at all locations in the study. In order to harmonize scans for all subjects in this real-world setting, we propose to use *mica* to estimate warping functions for the subset of subjects who have multiple scans, average these warping functions across subjects; and use the resulting mean to harmonize images for subjects with only a single scan available. We assessed the performance of this approach using leave-one-scan-out cross validation in the trio2prisma data. Specifically, we removed the Prisma scan for one subject and computed the *mica* warping functions 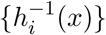 for the remaining subjects. We then computed the pointwise mean of these warping functions; using this as the warping function for the removed subject, we obtained a predicted Prisma scan from the known Trio scan. This process was repeated for each of the ten subjects. In the subsequent sections, scans harmonized using this leave-one-scan-out (*loso*) approach will be referred to below as *loso*-harmonized images and Trio scans harmonized using the full data will be referred to as *mica*-harmonized scans.

### 2.3 Statistical performance

All analyses were performed in the R software environment.

#### 2.3.1 NAIMS data

To assess the performance of our method on the NAIMS data we quantified T2-hyperintense lesion volume from the 3D FLAIR and T1-w images in both the White Stripe normalized and *mica*-harmonized images using MIMoSA (Valcarcel et al., 2018) for automated lesion segmentation. Because the number and volume of lesions are important metrics for monitoring MS disease progression (Bakshi et al., 2008) and the evaluation of therapeutic efficacy (Filippi et al., 2006), eliminating non-biological variability in detected lesion volumes will help clinicians deliver the best possible care to their patients.

We quantified mean and variance of lesion volumes within and across sites after applying White Stripe alone and after applying *mica* followed by White Stripe.

#### 2.3.2 trio2prisma data

For the trio2prisma data, we compared *mica* and *loso* to the histogram matching algorithm proposed by Nyúl et al. (1999), as implemented in Fortin et al. (2016). For better performance we first removed background voxels before running the histogram matching algorithm. To quantify performance of the methods we computed Hellinger distance of images before and after normalization, both within and across subjects. The Hellinger distance operates on PDFs of intensities, and its square is given by

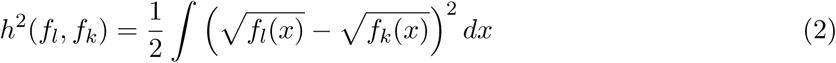

for PDFs *f*_*l*_(*x*) and *f*_*k*_(*x*). We visualized CDFs and PDFs and calculated Hellinger distances (Figures 4, 5, and 6, respectively) using images that had been *mica* or *loso*-harmonized but not yet White Stripe normalized. This is to isolate and visualize the effects of our method. For downstream analyses, including automated white and gray matter segmentation, we applied White Stripe normalization to the *mica* and *loso*-harmonized images to remove any residual intensity unit effects. We then estimated gray and white matter volumes and compare these across harmonization methods.

**Figure 4:**
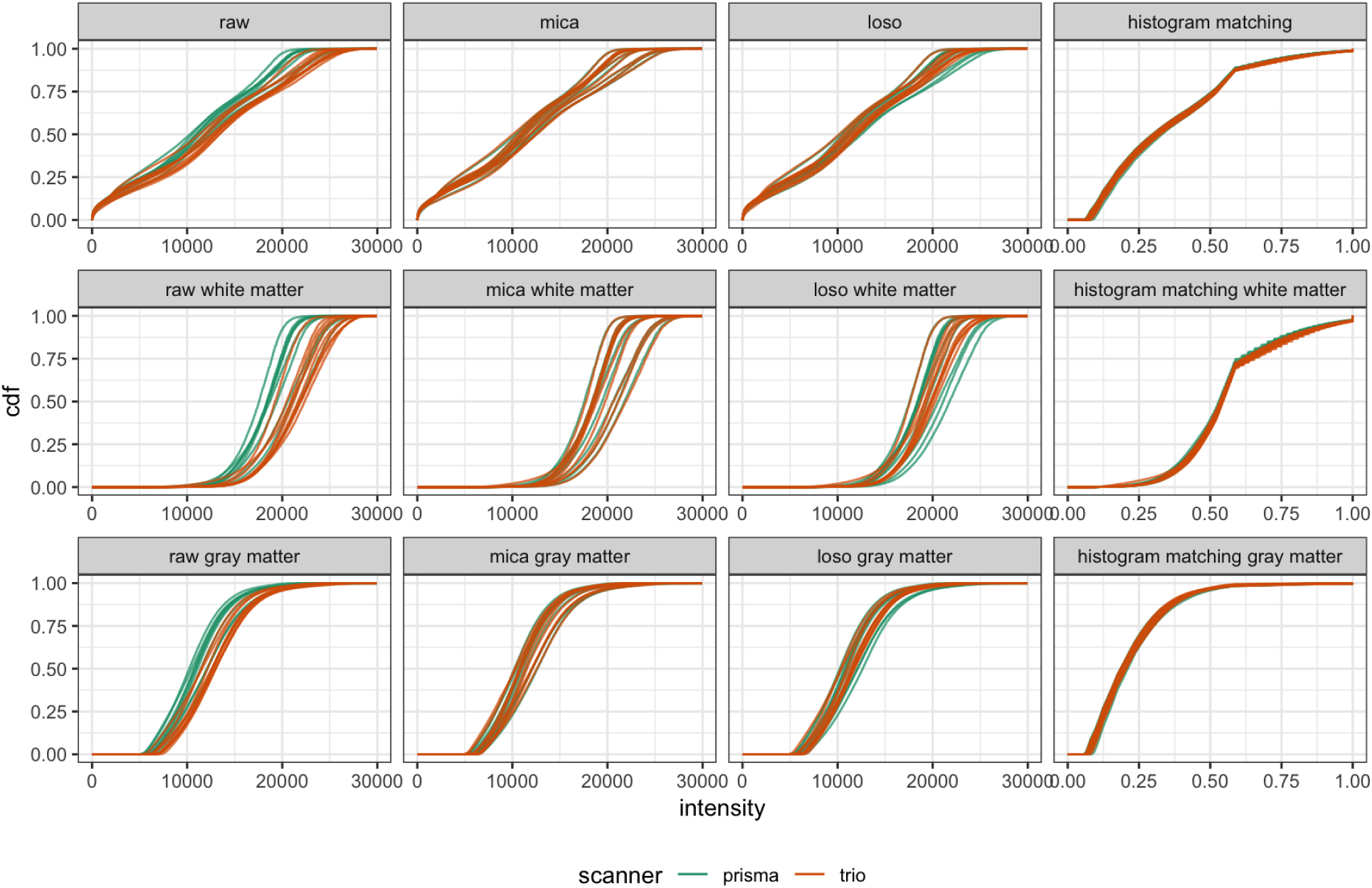
CDFs of intensities before and after harmonization by tissue type in the trio2prisma study. Rows indicate tissue type, with whole brain, white matter, and gray matter shown in rows 1, 2, and 3, respectively. Columns correspond to different harmonization methods.

**Figure 5:**
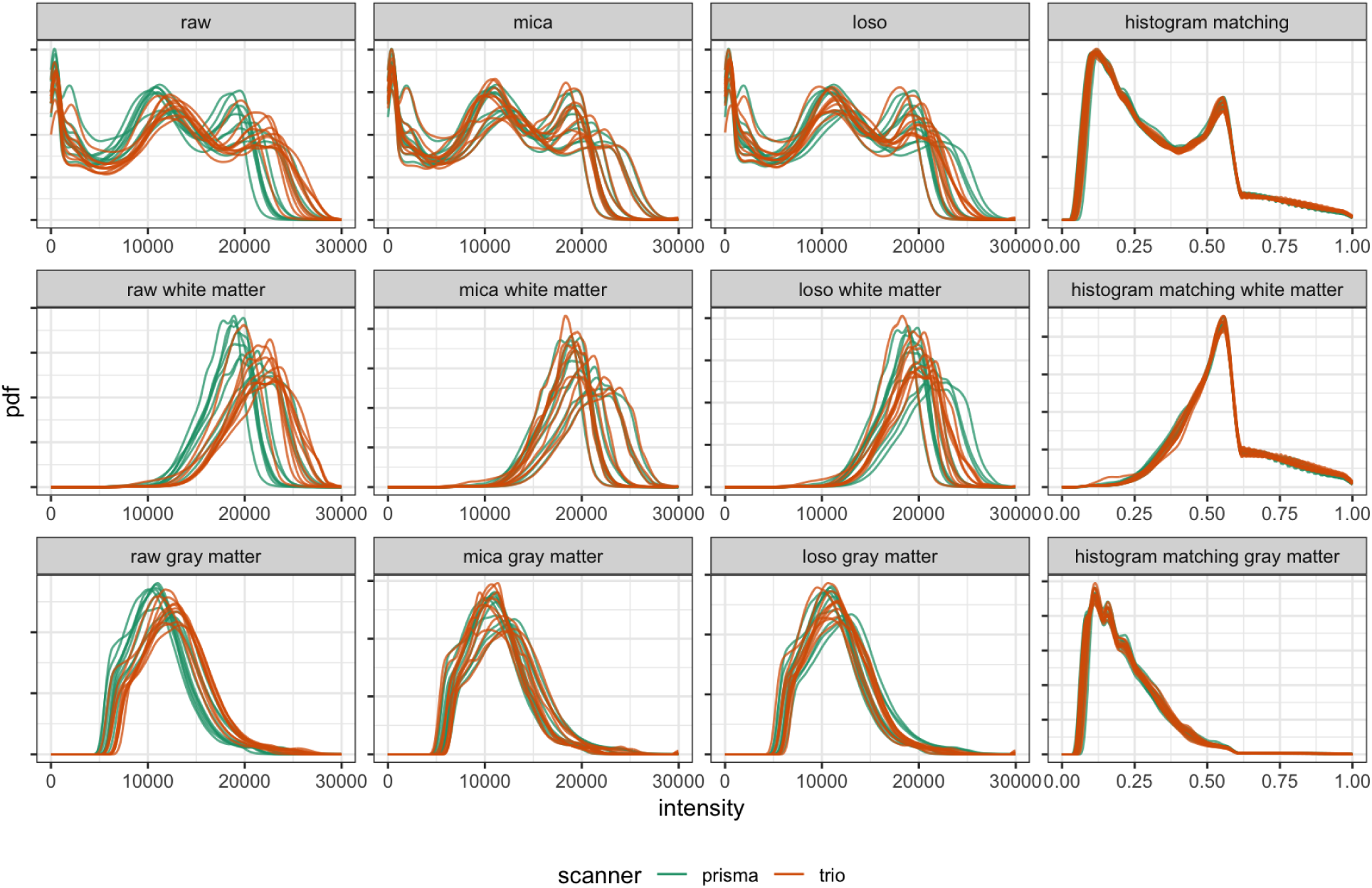
Histograms of intensities before and after harmonization by tissue type in the trio2prisma study. Rows indicate tissue type, with whole brain, white matter, and gray matter shown in rows 1, 2, and 3, respectively. Columns correspond to different harmonization methods.

**Figure 6:**
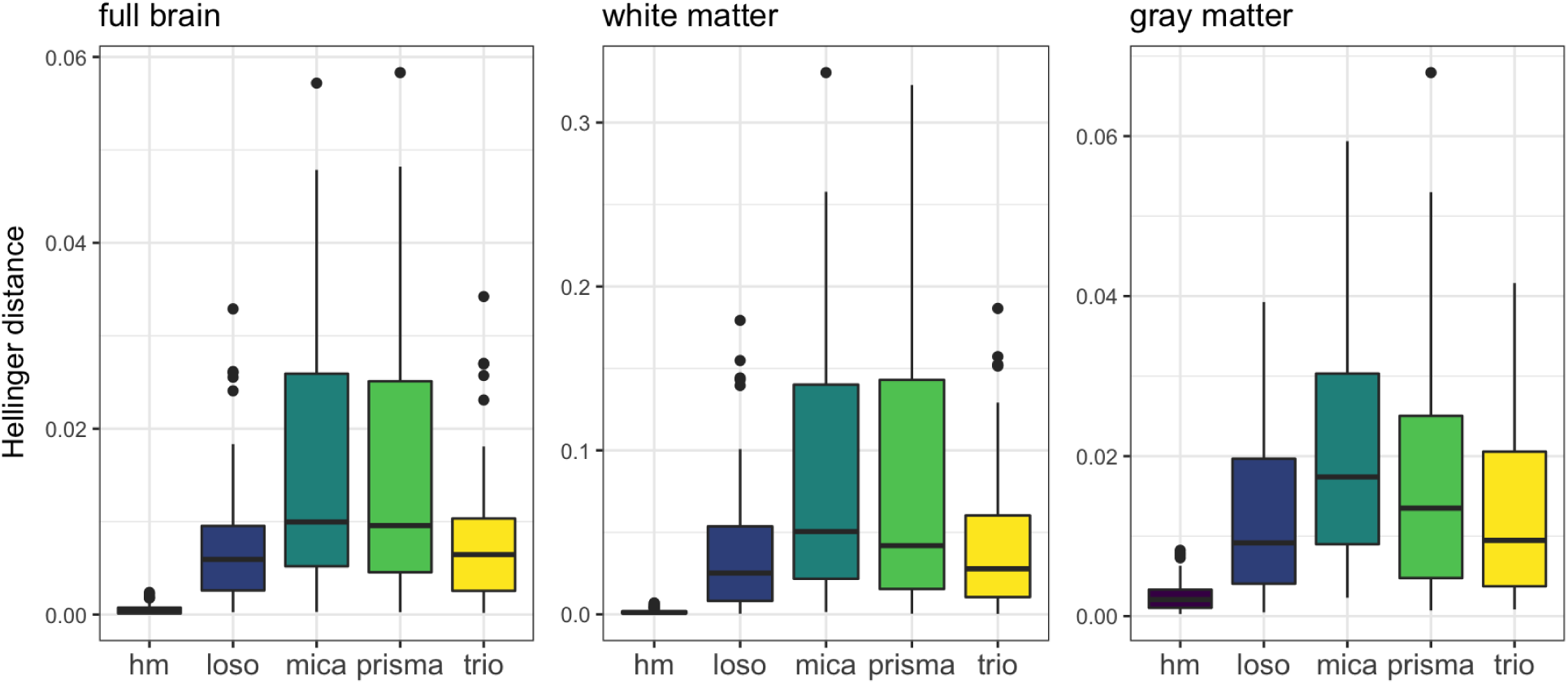
Boxplots of Hellinger distances across subjects, shaded by method. Columns show results for full brain (left), white matter (middle), and gray matter (right).

## 3 Results

For the NAIMS pilot data, we compared White Stripe normalized images to images processed using the *mica* approach outlined in section (2). For the trio2prisma data, we compared four harmonization strategies: no harmonization, histogram matching, *mica*, and *loso*. The main findings from these comparisons are summarized in the following two sections.

### 3.1 *mica* reduces variation in lesion volumes across sites in the NAIMS study

We *mica*-harmonized then White Stripe normalized the NAIMS scans, and then quantified MS lesion volume to assess the effect of scanner variability on a common downstream analysis before and after *mica* harmonization. The left panel of Figure 1 shows PDFs of raw voxel intensities from the NAIMS study images, and the right panel shows PDFs of images that have been *mica*-harmonized then White Stripe normalized. The raw PDFs show small differences within site, which are attributable to intensity unit effects, and larger differences across site, which are attributable to scanner effects. Scanner effects are particularly large between the UCSF site and other sites. After *mica* harmonization, the images across and within site have the same distributions of voxel intensities.

Figure 3 shows estimated T2-hyperintense lesion volume across sites for both White Stripe alone and White Stripe in conjunction with *mica* for scan-rescan pairs across the seven NAIMS sites. Compared to White Stripe alone, *mica* in conjunction with White Stripe yielded less variable lesion volume measurements across sites (variance 11.8 *ml*^2^ vs. 17.1 *ml*^2^) and similar lesion volume measurements within sites (variance 12.4 *ml*^2^ vs. 11.9 *ml*^2^). We see a larger impact across sites than within sites, suggesting that our method decreases site-to-site variance as expected and, together with White Stripe, performs comparably to existing methods for within site variance.

**Figure 3:**
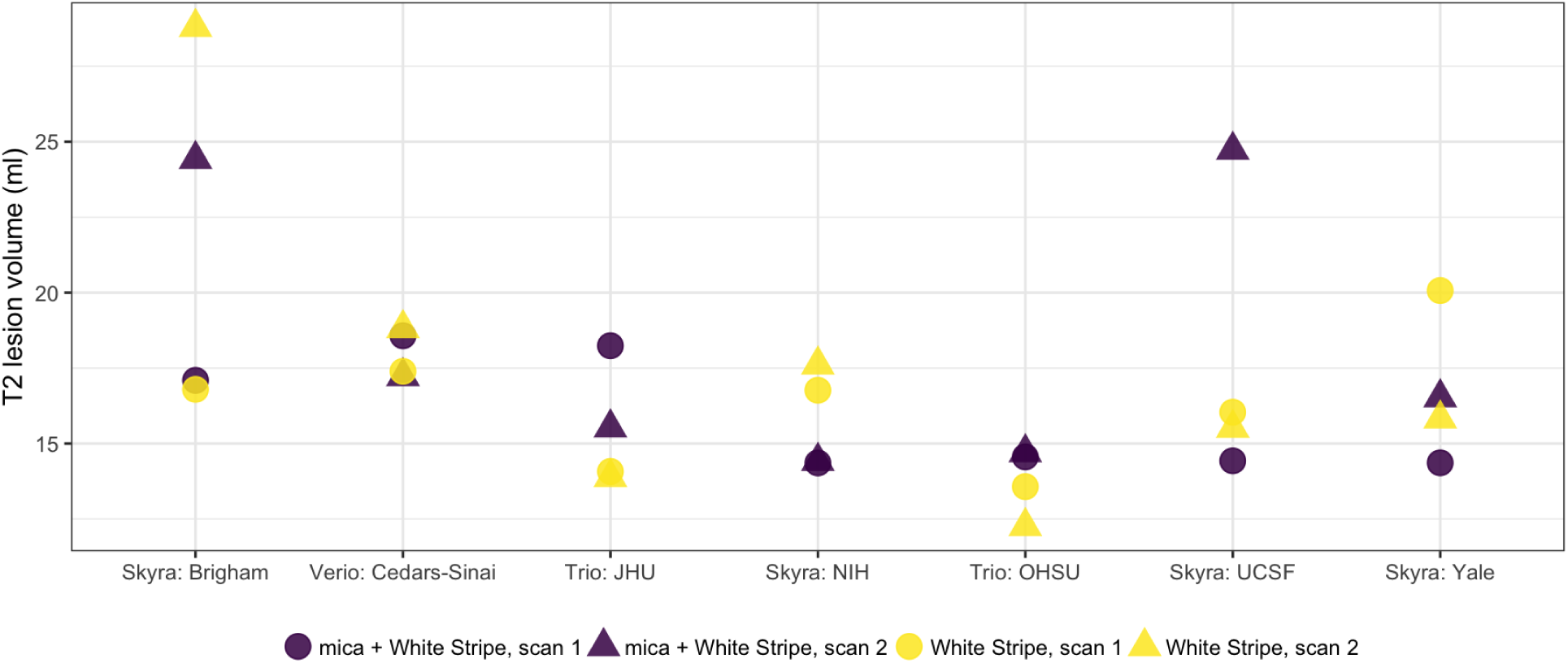
Estimated T2 lesion volumes for scan-rescan pairs at each of 7 sites in the NAIMS study. Circles indicate scan 1 and triangles indicate scan 2. Light and dark colors are volumes for White Stripe normalized images and *mica* normalized images, respectively.

### 3.2 *mica* preserves variation across subjects in the trio2prisma study

An appropriate harmonization method for multisite studies should reduce variability across scanners within the same subject but preserve biological differences across subjects. Here, we evaluate results from the trio2prisma data with these goals in mind. We compared *mica* and *loso*-harmonized images to images processed by histogram matching.

Figures 4 and 5 show CDFs and PDFs, respectively, under different harmonization scenarios. Visual inspection of intensity PDFs and CDFs in untransformed images suggests differences across scanners: the Prisma scans tend to have lower intensity values and higher peaks than the Trio scans. For both *mica* and histogram matching, within-subject technical variability is reduced because PDFs of Trio scans and Prisma scans are aligned. *mica* accomplishes this by mapping the Trio scan to the original Prisma scan, thus preserving the original features of the Prisma scans including variability across subjects. Histogram matching must be applied to scans from both the Trio and Prisma scanners, and reduces within-subject variability at the expense of eliminating desired differences across subjects. *loso* provides reasonable harmonization in that it maps Trio scans into the same range of intensity values as Prisma scans, but has less accuracy in reducing within-subject variability than *mica* or histogram matching. However, much of the desired across-subject variability is retained.

We quantified the variability across subjects using the Hellinger distance from equation (2) on PDFs of voxel intensities. Figure 6 displays boxplots of these pairwise distances for the original Trio scans, original Prisma scans, and scans processed by histogram matching, *loso*, and *mica*. The figure is divided into distances calculated on the full skull-stripped images (left column), white matter (middle column), and gray matter (right column). The *mica*-harmonized Trio scans have similar across-subject variability to the Prisma scans. The *loso* scans have variability comparable to the original Trio scans but smaller than the Prisma scans. Histogram matching virtually eliminates inter-subject variability, including that which is presumably biological.

Figure 7 shows an axial slice of the Trio image for one subject from the trio2prisma dataset. The slice is shown for raw intensity values (center), intensity values after *mica* harmonization (left), and intensity values after histogram matching (right). Here, *mica*-harmonization brightens the contrast between white and gray matter but does not distort the shape of biological features in the tissue. Histogram matching, however, drastically changes the appearance of the image, converting some gray matter to CSF and some white matter to gray matter.

**Figure 7:**
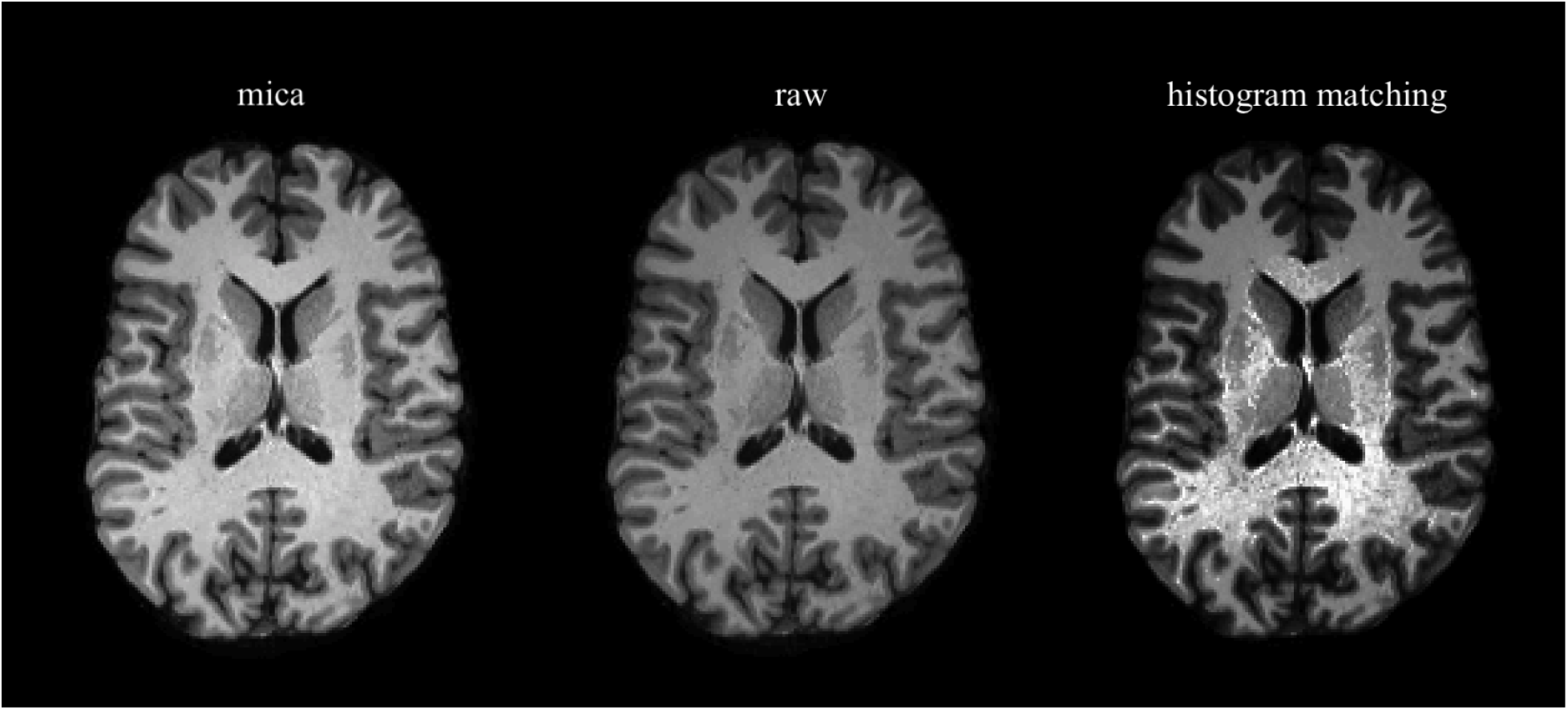
Axial slice of skull-stripped images from a single subject in the trio2prisma dataset. Center panel shows the raw intensity values from an image collected on the Trio scanner. Left and right panels show the same image after *mica* harmonization and histogram matching, respectively.

Finally, neither harmonization nor normalization methods should bias assignment of tissue type. After harmonization or normalization, we expect that segmentation volumes from harmonized Trio scans should be similar to segmentation volumes from unharmonized and unnormalized (raw) Trio scans. We estimated white and gray matter volumes on original Trio scans and after histogram matching, White Stripe, *mica* followed by White Stripe, and *loso* followed by White Stripe. Figure 8 shows these volumes for each subject and tissue type. All methods have at least some difference in segmentation volume compared to the raw data. The *mica, loso*, and White Stripe methods all performed similarly, with volumes that are close to those of the raw images but slightly lower for the gray matter and slightly higher for the white matter. Histogram matching, however, had much lower segmentation volumes in both the gray matter and the white matter than either the raw data or any other method. As shown in Figure 7, histogram matching severely distorts the image; we believe this distortion causes the segmentation algorithm to convert some gray matter to CSF and some white matter to gray matter, which explains the consistently lower volumes.

**Figure 8:**
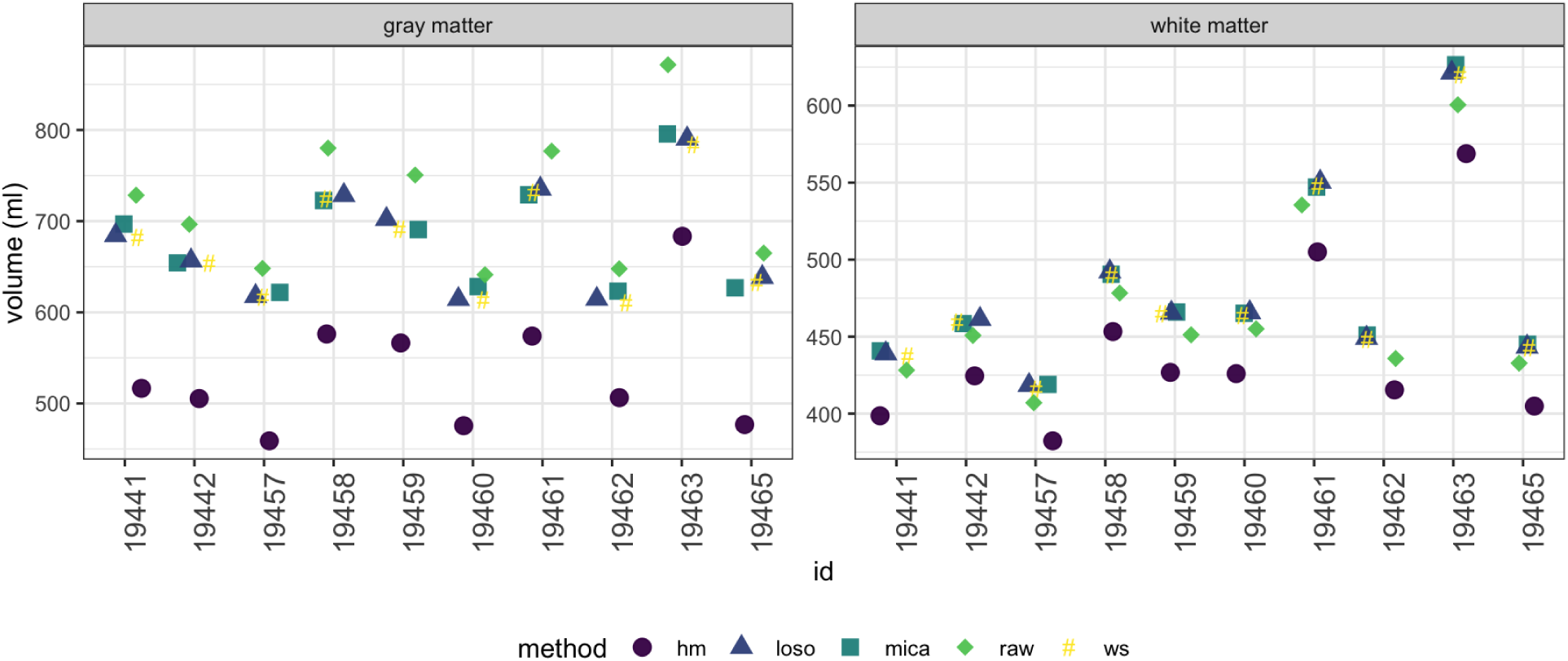
Segmented brain volume in the gray matter (left) and white matter (right) for each trio2prisma subject across harmonization approaches. We compare no normalization or harmonization (raw), histogram matching (hm), White Stripe normalization (ws), *mica*, and *loso*.

### Discussion

Unwanted technical variability due to scanner effects in multisite clinical trials and observational studies is an increasingly common problem; to mitigate these scanner effects we introdce *mica*, a method that harmonizes structural MRI images by defining nonlinear transformations between CDFs of voxel intensities. To specifically target scanner effects, we developed a paradigm for understanding scanner effects and intensity unit effects as related but distinct sources of technical variability in MRI scans. Intensity unit effects are due to arbitrary MRI unit intensities within a single scanner, and scanner effects are unwanted technical artifacts introduced across scanners or sites. We also distinguish between approaches targeting these sources of variability: normalization methods address intensity unit effects, and harmonization methods, the focus of our study, address scanner effects.

Our data came from two small studies, the NAIMS pilot study and the trio2prisma study, with multiple images per subject taken on multiple scanners, and nonlinear scanner effects. We found that *mica* reduced within-subject variability in whole brain scans as well as white and gray matter while preserving biological variability across subjects. We also found that *mica*, paired with White Stripe, enhanced reproducibility of measurements of MS lesion volume across sites.

Normalization methods such as histogram matching and White Stripe are sometimes used for harmonization, but they are inadequate in cases where across-site differences are much larger than those within site. Additionally, histogram matching can reduce biological variability across subjects and White Stripe can leave residual technical variability in the gray matter. While we differentiate conceptually between intensity unit effects and scanner effects, we also acknowledge that in reality these artifacts can be challenging to separate. As a result, *mica* is likely to remove some intensity unit effects and intensity normalization methods are likely to remove some scanner effects when applied separately. In particular, White Stripe alone will likely perform well as a harmonization method when scanner effects are small, linear transformations. Histogram matching, however, is likely to remove desired variability across subjects and bias results.

Because our method is flexible and operates on the full brain, we can map images from one scanner to another. This mapping is only exact for a particular subject when images are available from both scanners, which is not realistic for most studies. That said, our leave-one-scan-out analysis suggests that when systematic site differences are present, *mica* can help understand scanner effects and mitigate those differences. Before conducting multisite studies, we recommend obtaining a baseline measurement of scanner variability by having a subset of patients measured at all sites. Our method can then be applied to all images collected to remove average scanner variability. We acknowledge that this solution is imperfect in the sense that average scanner variability collected from a subset of patients in a trial will not always capture the true scanner variability for each subject. However, our simple and easy-to-apply methodology is an important step forward for an increasingly prevalent problem. There is evidence that scanner effects may vary across covariates such as gender and age, so extensions to *mica* that incorporate covariates may address some of the issues outlined above.

### 5 Software

To enable use of *mica* we have written an *R* software package which is available for download at https://github.com/julia-wrobel/mica.

## Acknowledgements

This work was supported by the National Institutes of Health R01NS085211, R21NS093349, R01MH112847, R01NS060910, R01EB017255, R01HL123407, R01NS097423, and the National Multiple Sclerosis Society RG-1707-28586, and the Race to Erase MS Foundation. The content is solely the responsibility of the authors and does not necessarily represent the official views of the funding agencies.

